# Impaired oxytocin signaling in the central amygdala in rats with chronic heart failure

**DOI:** 10.1101/2023.11.22.568271

**Authors:** Ferdinand Althammer, Ranjan K. Roy, Matthew K. Kirchner, Elba Campos Lira, Stephanie Schimmer, Alexandre Charlet, Valery Grinevich, Javier E. Stern

## Abstract

**Aims:** Heart failure (HF) patients often suffer from cognitive decline, depression, and mood impairments, but the molecular signals and brain circuits underlying these effects remain elusive. The hypothalamic neuropeptide oxytocin (OT) is critically involved in the regulation of mood, and OTergic signaling in the central amygdala (CeA) is a key mechanism controlling emotional responses including anxiety-like behaviors. Based on this, we used in this study a well-established ischemic rat HF model and aimed to study alterations in the hypothalamus-to-CeA OTergic circuit.

**Methods and Results:** To study potential HF-induced changes in the hypothalamus-to-CeA OTertic circuit, we combined patch-clamp electrophysiology, immunohistochemical analysis, RNAScope assessment of OTR mRNA, brain region-specific stereotaxic injections of viral vectors and retrograde tracing, optogenetic stimulation and OT biosensors in the ischemic HF model. We found that most of OTergic innervation of the central amygdala (CeA) originated from the hypothalamic supraoptic nucleus (SON). While no differences in the numbers of SON➜CeA OTertic neurons (or their OT content) was observed between sham and HF rats, we did observe a blunted content and release of OT from axonal terminals within the CeA. Moreover, we report downregulation of neuronal and astrocytic OT receptors, and impaired OTR-driven GABAergic synaptic activity within the CeA microcircuit of rats with HF.

**Conclusions:** Our study provides first evidence that HF rats display various perturbations in the hypothalamus-to-amygdala OTergic circuit, and lays the foundation for future translational studies targeting either the OT system or GABAergic amygdala GABA microcircuit to ameliorate depression or mood impairments in rats or patients with chronic HF.

## Introduction

Cardiovascular diseases are a major cause of morbidity and mortality worldwide, and heart failure (HF) is a growing public health concern. It is estimated that over 64 million people worldwide suffer from heart failure, and this number is expected to rise in the coming years ^1^. Numerous studies in the past decades have focused on identifying genetic and lifestyle risk factors for HF, as well as peripheral and central consequences of myocardial infarction ^2–5^. Conversely, much less is known about HF-associated impairments in cognitive function and emotional control, and the underlying brain circuits involved. Given the high incidence of depression ^6–9^, anxiety^8,10–12^ and cognitive impairment in HF patients ^13–15^, a detailed understanding of the underlying mechanisms is paramount for the development of adequate therapeutic interventions.

Different HF-based studies and methodological approaches revealed numerous mechanisms and signals that contribute to neurohumoral and cardiovascular alterations in HF ^16,17^. Among the identified molecules and potential contributors, the hypothalamic neuropeptide oxytocin (OT), traditionally known for its role in social and reproductive behavior, has recently been implicated in the onset and pathophysiology of HF ^18–20^. For example, recent work from our group has shown increased excitability of magnocellular neurosecretory OT neurons in rats with HF ^21,22^. Moreover, activation of medullar-projecting hypothalamic OT neurons have been shown to contribute to cardiac sympathetic nerve activation and cardiac arrhythmias following acute myocardial infarction in rats ^23,24^. Within the medulla, OT also modulates parasympathetic outflow ^25^, and a recent study demonstrated that chronic activation of OT neurons improved cardiac function during the chronic stages of HF ^26^. Thus, OT has been shown to play pleiotropic roles in autonomic regulation of cardiac functions both in the acute and chronic phases of HF. Still question remain whether OT contributes also to cognitive/mood disorders in HF, and if so, what the cellular mechanisms and neuronal substrates are.

The central nucleus of amygdala (CeA) is critical for the processing of emotional and social cues ^27–29^ and is composed of distinct subnuclei, including the central medial (CeM) and central lateral amygdala (CeL). Various studies highlighted that OT release within the CeA is critical for the acquisition, expression and extinction of contextual fear, as well as for the modulation of emotional components of fear experiences in both rats and humans ^30–37^. Recent studies have shown that within the CeA, OT release from hypothalamic inputs activates oxytocin receptor (OTR)-expressing GABAergic interneurons located in the CeL. This effect resulted in an increased GABAergic inhibition of neurons located in the CeM, decreasing their firing output to the brainstem. Thus, OT indirectly inhibited GABAergic CeM neurons, supporting the idea that OT directly regulates the activity of local inhibitory CeA circuits ^38^. The functional implication of these actions is further supported by our previous work showing that OT activation of GABAergic CeL interneurons mediates a powerful anxiolytic effect ^33^ and that OT release within the CeL is necessary for the extinction of fearful memories ^31^. More recently, we have demonstrated that not only neurons, but also OTR-expressing astrocytes contribute to OT actions within the CeL→CeM circuit, influencing neuronal activity via activity-dependent release of D-serine ^30^.

Taken together, these studies support a major role of OT in the CeA GABAergic synaptic activity modulating normal emotional responses and mood. Still, whether changes in OT signaling within the CeA occurs in rats with HF failure, and whether this constitutes a potential mechanism contributing to mood disorders in this disease, remains unknown. To address this question, we used a well-established rat myocardial-infarction HF model ^10,21,39–43^, in which we have previously identified signals and mechanisms contributing to neurogenic component of humoral and cardiac components of this disease ^10,21,39–41^. Importantly, we previously demonstrated an impaired mood and cognitive phenotype in this rat model including increased anxiety and anhedonia ^10,39^, and provided evidence for the CeA as a potential neuronal substrate mediating these effects ^10^. Based on this evidence, we aimed here to investigate the impact of HF on hypothalamic CeA-projecting OT neurons, OTR mRNA and protein levels, as well as the OT-mediated functional connectivity of the local CeL→CeM circuit. Thus, we utilized patch-clamp electrophysiology, immunohistochemical analysis, RNAScope assessment of OTR mRNA, brain region-specific stereotaxic injections of viral vectors and retrograde tracing, optogenetic stimulation and OT biosensors in the rat ischemic HF model to study these possibilities. In our HF rats, we observed depletion of OT from OTergic axon terminals within the CeA, alterations in OT trafficking and release, a reduction of neuronal and astrocytic OTRs and perturbations in the OT-sensitive, neuronal GABAergic CeL→CeM circuit, which potentially underlie the increased fear and anxiety in rats and patients with chronic heart failure.

## Results

### The hypothalamic SON is a major source of OT innervation to the CeL in male rats

It has previously been demonstrated that the SON is the major source of OTergic innervation to the CeL in female Wistar rats ^31,33^. To validate whether this is also the case in male Wistar rats, we injected rats with OT cell type specific recombinant adeno-associated virus (AAV_OT_p_-Venus) into the SON and PVN, two of the major OT outputs in the brain (n=6 rats, left hemisphere SON, right hemisphere PVN) (**Figure 1A**) and subsequently analyzed OT fiber density within the CeA on the ipsilateral side of each injection. As show in in **Figure 1B, C** we found that the SON provided a significantly denser OT innervation to the CeL compared to the PVN. To rule out a significant contribution of contralateral CeL-projecting fibers originating from the PVN, we injected 2 rats unilaterally into the PVN and analyzed the contralateral CeL (not shown). Here, we observed only sparse OT fibers, a finding that is in line with the previously reported data on female Wistar rats ^33^. Thus, we concluded that similar to females, the ipsilateral SON is the primary source of OT innervation to the CeL in male rats, and thus exclusively focused on the SON_OT_-CeL pathway.

**Figure 1.**
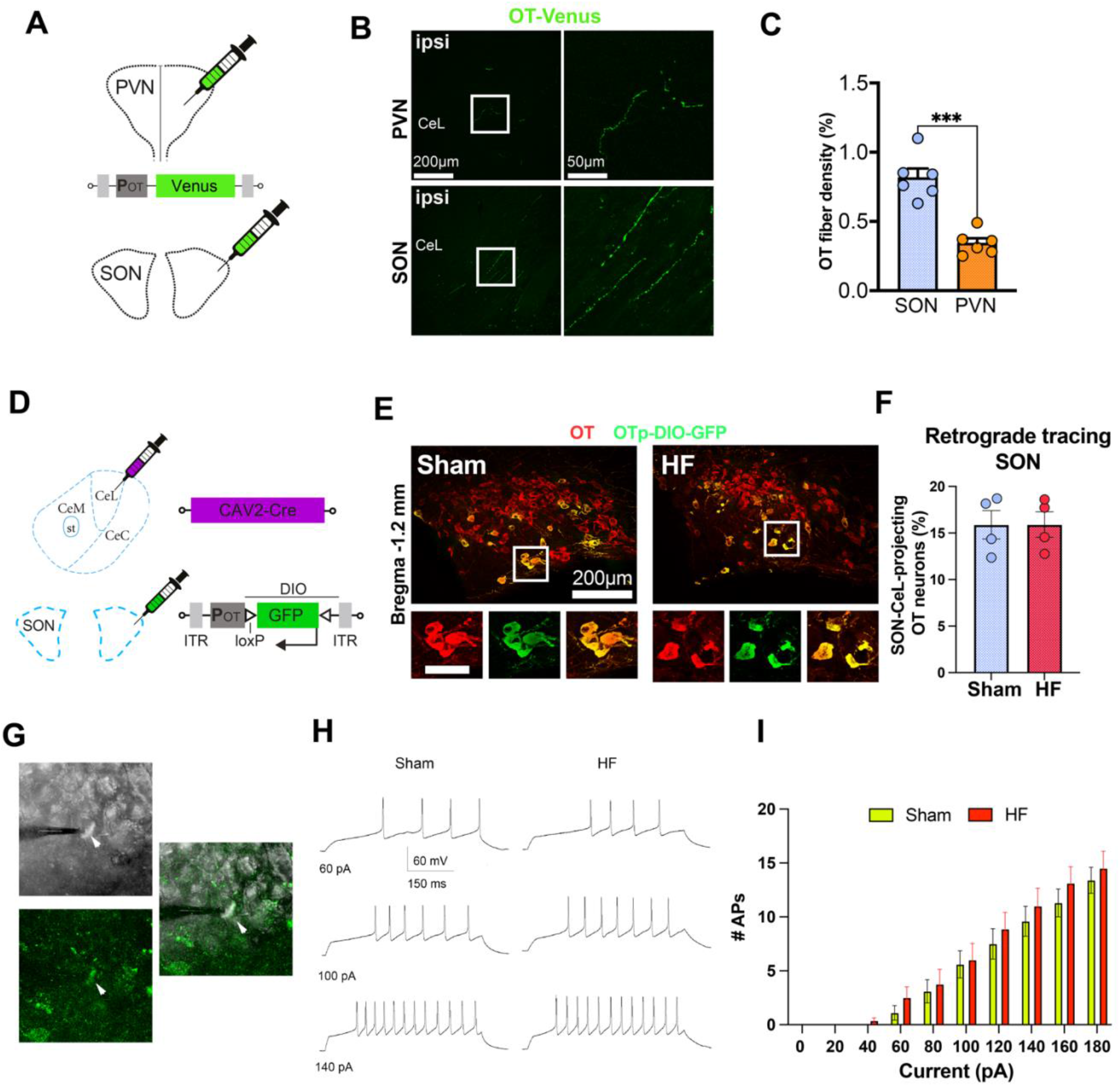
Anatomical and functional assessment of CeA-projecting SON OT neurons. **A** Schematic depiction of viral injections into hypothalamic nuclei. AAV-OT_p_-Venus was injected unilaterally into PVN and SON of male WT rats (n=6). **B** Representative confocal images showing OTergic innervation of PVN and SON after 4 weeks of unilateral viral infection and OT_p_-driven Venus expression. **C** Quantification of OT fibers density in the SON and PVN following viral injections, ***p <0.001; Mann-Whitney test. **D** Schematic depiction of viral approach to retrogradely label CeL-projecting SON OT neurons. WT and HF rats received bilateral injections of CAV2-Cre into the CeL and AAV_OT_p_-DIO-GFP into the SON. **E** Representative confocal images of retrogradely-labeled, CeL-projecting SON OT neurons in WT and HF rats. Immunohistochemical labeling via the PS38 antibody is shown in red, intrinsic, virally-expressed GFP in green, overlay in yellow. Enlarged insets depict co-localization of red and green fluorescent signals. **F** Quantification of the number of CeL-projecting SON OT neurons in WT and HF rats (n=4). **G** Representative sample of a patched SON_OT_-CeL-projecting neurons identified by the presence of intrinsically expressed GFP fluorescence (green). Images obtain under DIC illumination (upper), fluorescence illumination (lower) and their overlap (right) images of the patched neuron is shown. **H** Samples of evoked spiking in SON_OT_-CeL-projecting neurons obtained from a Sham and HF rat following evoked by 60, 100 and 140 pA depolarizing pulses are shown. **I** Plot of the mean number of evoked action potentials (APs) as a function of variable current injections for SON_OT_-CeL-projecting neurons obtained from a Sham (n= 4) and HF (n= 5) rats.

### SON_OT_-CeL-projecting neuron numbers or excitability is not altered in rats with HF

We next asked whether the numbers and/or electrophysiological properties of SON_OT_-CeL projecting neurons were altered in rats with HF (cardiac parameters assessed via echocardiography can be found in **Figure S1**). To this end, rats were injected with CAV2-Cre ^31,44,45^ into the CeL and AAV_OT_p_-DIO-GFP into the SON (**Figure 1D**). This approach allowed us to selectively label SON OT neurons projecting to the CeL. Representative examples are shown in **Figure 1E**. This complementary approach further confirms the SON as a source of OT innervation of the CeL, and indicated that approximately 15% of SON OT neurons provide innervation to the CeL, a proportion that was not different from that observed in HF rats (**Figure 1F**).

To determine whether the intrinsic excitability and repetitive firing properties of SON_OT_-CeL-projecting neurons were altered in HF, we used acute *ex vivo* slices from rats that received the double injection protocol described above, to whole-cell patch clamp recordings from identified SON_OT_-CeL-projecting neurons in sham and HF rats (**Figure 1G-I**). For this, neurons were held at a hyperpolarized membrane potential (Vm = –80 mV) and subjected to depolarizing pulses of increasing magnitude. A plot of the number of evoked action potentials (APs) per current pulse was built (e.g., input-output function (I/O)) for both sham and HF rats. As depicted in **Figure 1I**, our results showed no differences in the I/O function between sham and HF rats (2-way ANOVA, F= 0.44, P= 0.52, n=10 and 8 in sham and HF rats, respectively). In addition, no differences in mean input resistance were observed between the two groups (sham: 540.1 ± 60.8 MΩ, HF: 616.2 ± 53.7 MΩ, unpaired t test, p= 0.36).

Taken together, our data suggest that neither the numbers nor the overall excitability and repetitive firing properties of SON_OT_-CeL-projecting neurons were affected in rats with HF.

### Reduced SON_OT_-CeL innervation and OT axonal release in rats with HF

To determine whether the degree of OT innervation of the CeL was altered in HF rats, we stained CeL-containing brain sections of sham and HF rats with an OT antibody (PS38, Neurophysin-I) (**Figure 2A-C**). Intriguingly, we found a significant reduction of OT immunoreactivity in fibers located within the CeL of HF rats (**Figure 2B, C**). However, when we assessed OT fiber densities following injection of AAV_OT_p_-Venus into the SON (as in **Figure 1A**), a similar degree of OT fiber densities were observed between the two groups (**Figure 2B, C**). These findings suggest a reduction in OT content in SON_OT_-CeL axonal terminals, rather than a decrease in the density of OT innervation in the CeL of HF rats.

**Figure 2.**
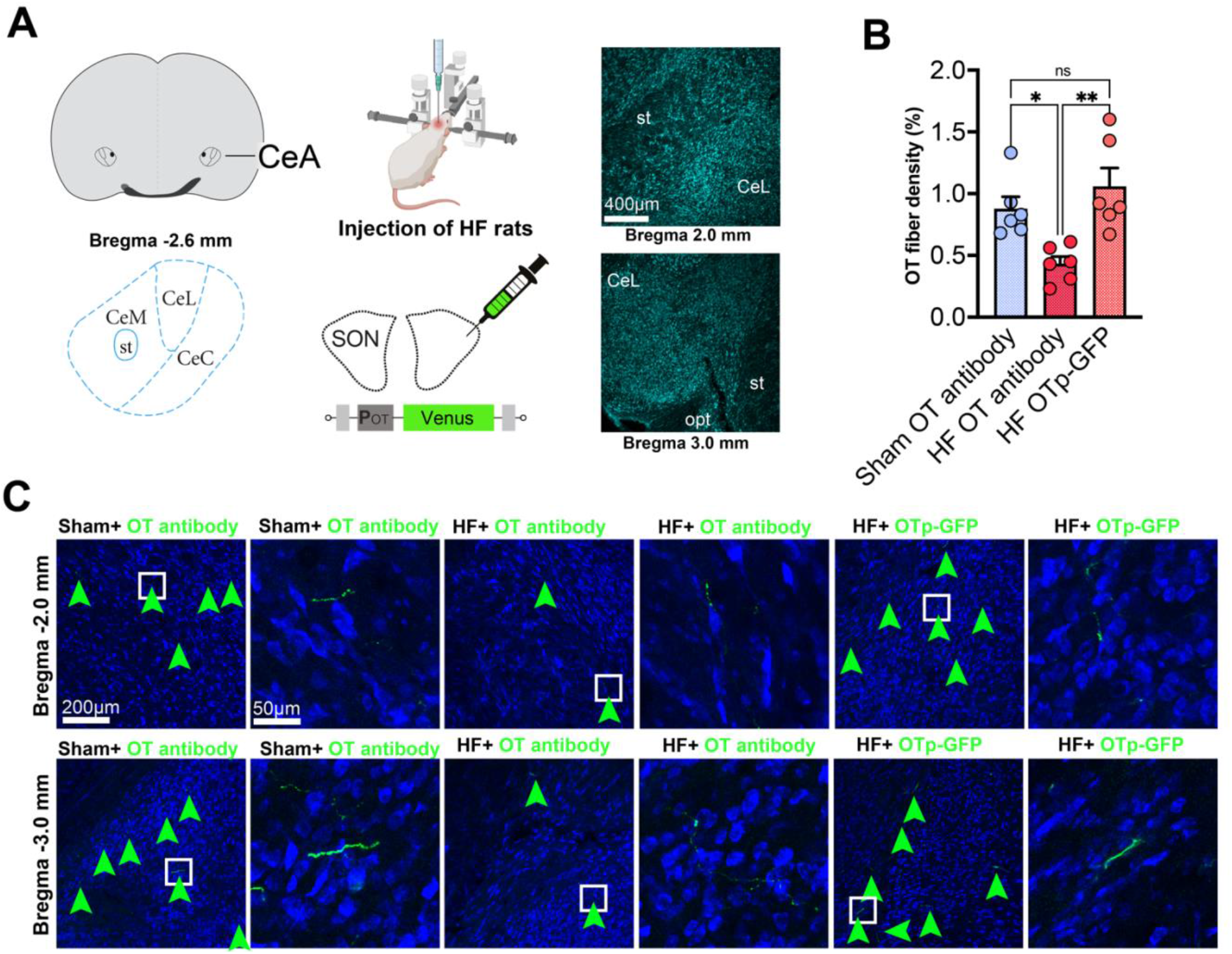
Assessment of OTergic fibers within the CeA in HF rats. **A** Anatomical location of CeL and CeM and schematic depiction of viral injections in HF rats. HF rats received bilateral injections of AAV_OT_p_-Venus into the SON. st=stria terminalis, opt=optic tract. **B** Quantification of OT fiber density in the CeL in sham rats stained via PS38, HF rats stained via PS38 and HF rats injected into the SON with AAV_OT_p_-Venus, *p <0.05; **p <0.01; Mann-Whitney test. **C** Confocal panel shows representative images of all three conditions quantified in **B** at two different Bregma levels. Green arrowheads indicate the presence of OTergic fibers within the CeL either labeled via PS38 in sham and HF rats or via AAV_OT_p_-Venus in HF rats. Enlarged insets show subtle differences in thickness and length of OTergic fibers within the CeL.

To further confirm a decreased OT-immunoreactive axonal content in the CeL of HF rats we used a complementary approach in which we directly measured activity-dependent OT release from SON_OT_-CeL-projecting fibers in sham and HF rats. To this end, we used CHO cells transfected with human OT receptor (Sniffer^OT^) and a red-shifted fluorescent Ca^2+^ indicator (R-GECO), an approach we recently show to detect endogenous release of neuropeptides with extremely high sensitivity and spatiotemporal resolutions ^46^. Sniffer^OT^ cells were platted into *ex vivo* slices containing the CeA that were obtained from sham and HF rats that received an injection of AAV_OT_p_-Chr2mCherry in the SON (**Figure 3A-B**). Laser stimulation of SON_OT_-CeL fibers expressing ChR2 in the CeL resulted in local release of OT that was efficiently detected by the surrounding Sniffer^OT^ cells (**Figure 3D,E**). While we found that the proportion of responsive Sniffer^OT^ cells was lower in slices derived from HF rats (19.7%) compared to sham controls (24.5%), this difference did not reach statistical significance (**Figure 3C**, p>0.05, chi-square test). In contrast, quantitative Ca^2+^ responses of Sniffer^OT^ differed significantly in HF compared to sham controls (**Figure 3E, F**). Indeed Sniffer^OT^ displayed longer response latency (p<0.01), shorter duration (p<0.05), and smaller integrated area Ca^2+^ responses (p<0.05) in HF compared to sham (**Figure 3F**). Peak Ca^2+^ amplitude was not different between the two groups (**Figure 3F**; p>0.05). Together, these results support a blunted activity-dependent release of OT from SON_OT_-CeL terminals in rats with HF.

**Figure 3.**
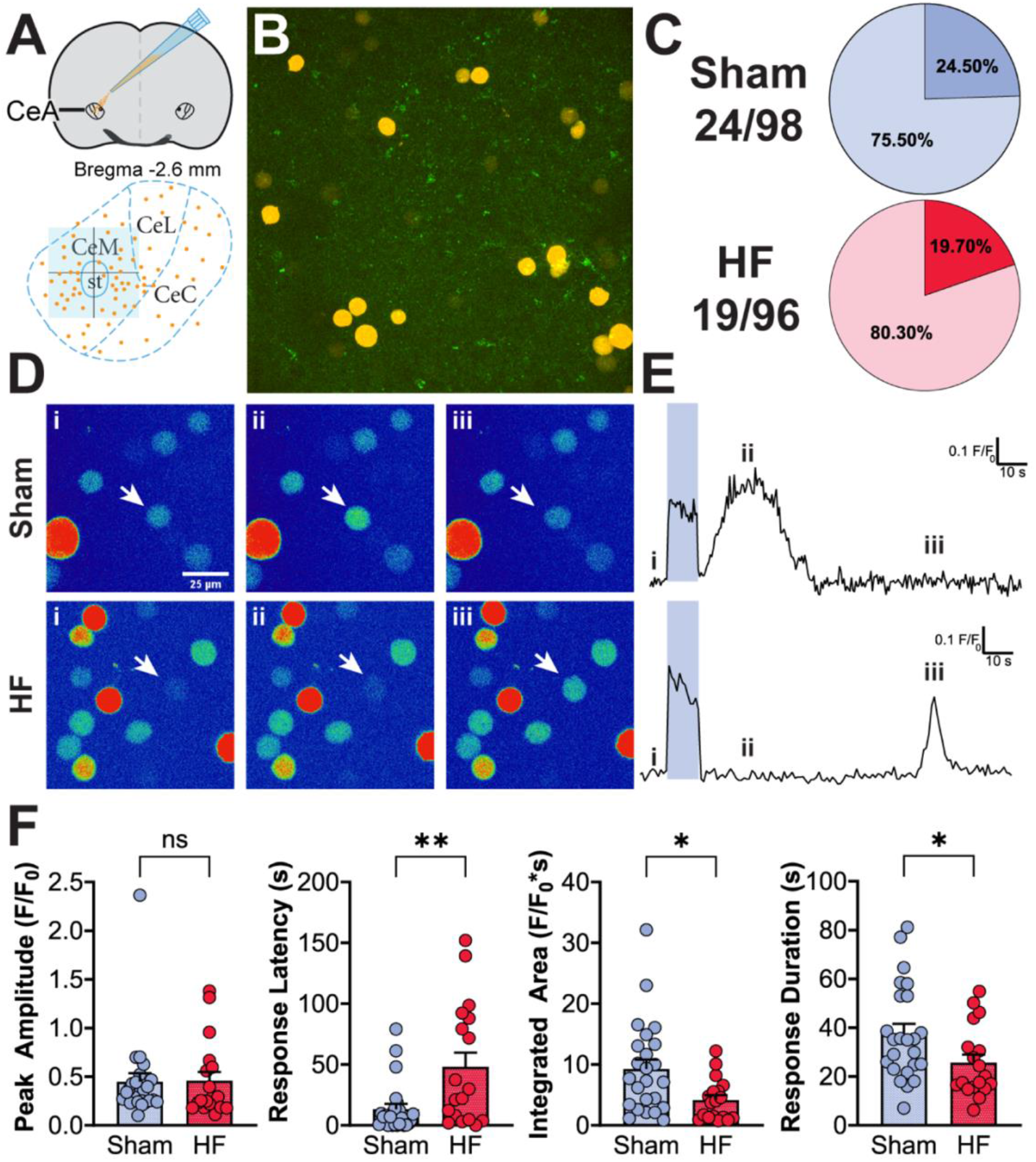
Impaired OT release from OT axonal terminals within the CeA of HF rats. **A** Anatomical location of sniffer cell transfer onto coronal brain slices containing CeL. Blue zone indicates region imaged. **B.** Fluorescence image of AAV_OT_p_-Chr2mCherry-infected terminals in CeM with Sniffer^OT^ seated on top of the slice. **C.** Response rates of Sniffer^OT^ to optogenetic Chr2 stimulation. **D.** Pseudocolor Ca^2+^ images showing responsive Sniffer^OT^ (arrow) at baseline (i), immediately after stimulation (ii), and much later after stimulation (iii) in a Sham (*top*) and HF (*bottom*) slice. **E.** Corresponding Ca^2+^ traces. Roman numeral timepoints correspond to images in (D). **F.** Comparison of quantified Ca^2+^ responses between sham and HF. Peak amplitude p>0.05, Response latency **p<0.01, integrated area *p<0.05, and response duration *p <0.05; Mann-Whitney test.

### Decreased OTR expression in the CeL of HF rats

We next wanted to determine whether in addition to changes in OT fiber content and release, there were also changes in the expression of OTRs within the CeA. We first evaluated OTR mRNA expression via qPCR using CeL-specific tissue punches as previously described ^10^. As shown in **Figure S2A**, we found a significant decrease in OTR mRNA in the CeL of HF rats relative to sham rats. We have previously shown that functional OTRs are expressed both in neurons and astrocytes in the CeA^30^. To address cell-type specific changes in OTR mRNA, we performed an approach combining simultaneously RNAScope hybridization against OTR mRNA with immunohistochemical staining against the astrocyte marker glutamine synthetase (GS) ^39,40,47^ (**Figure 4A,B**). Neurons were identified using our recently published approach that enables to differentiate neurons from glial cells based on DAPI size measurements (^39^, see also **Figure S2B**). Our quantitative analysis revealed a significant decrease in OTR mRNA signal in CeL neurons of HF rats, without changes in the proportion of CeL neurons expressing OTRs (**Figure 4C**). The opposite pattern was observed in CeL astrocytes, in which we found a decrease in the proportion (but not in the individual OTR content) of astrocytes expressing OTRs in HF rats compared to Sham rats. (**Figure 4D**). No differences in the number of neurons or astrocytes within the CeL were observed between sham and HF rats (**Figure S2C**).

**Figure 4.**
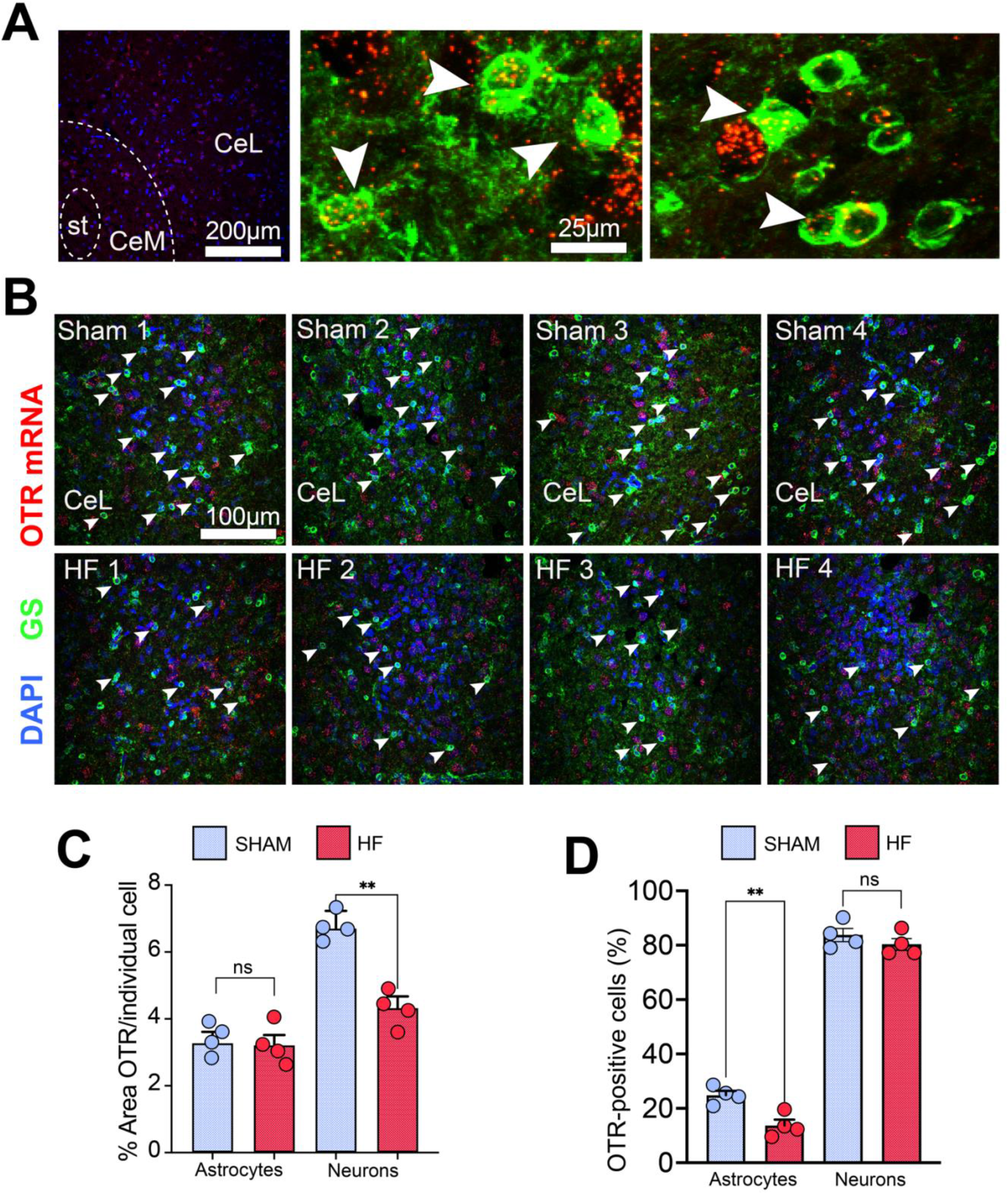
Alterations of neuronal and astrocytic OTR expression in the CeA of HF rats. **A** Overview image displays the distribution of OTR mRNA-positive cells within the CeL visualized via RNAScope hybridization. DAPI=blue, OTR mRNA=red, st=stria terminalis. High magnification images show OTR-positive astrocytes within the CeL labeled via the astrocyte marker glutamine synthetase (GS, green). White arrowheads indicate co-localization of OTRmRNA (red) and GS (green). **B** Confocal panel shows 4 RNAScope hybridization and GS labeling within the CeL of 4 different sham an HF rats. White arrowheads indicate OTR mRNA-positive astrocytes. **C** Cell type-specific quantification of OT mRNA signal (single cells, neurons vs. astrocytes) in sham and HF rats (n=4). **D** Cell type-specific assessment of the number of OTR-positive neurons and astrocytes in sham and HF rats (n=4), **p <0.01 Mann-Whitney test.

To confirm that the decrease OTR expression at mRNA level was also observed at the protein level, we evaluated overall OTR immunoreactivity in the CeL of sham and HF rats using an antibody that we recently validated^40^. As shown in **Figure S2D, E**, we observed a significant reduction of OTR protein immunoreactivity in the CeL of HF when compared to Sham rats.

Collectively, these observations suggest a blunted OT signaling efficacy and/or disturbed OT-mediated astrocyte-neuron communication with the CeA of HF rats.

### Blunted inhibitory effect of OT in CeM output neurons in rats with HF

Within the CeA, OTRs are highly localized in the CeL, where OT release stimulates increased firing activity of GABAergic interneurons, which suvsequently inhibits output projection neurons in the CeM (see **Figure 5A** ^33,38^). To determine whether the anatomical and molecular changes in the SON_OT_-CeL pathway in HF rats had functional implications and affected local circuitry activity within the CeA, we performed electrophysiological whole-cell patch recordings from neurons in the CeM from sham and HF rats that received an injection of AAV_OT_p_-Channelrhorodopsin2 (Chr2)mCherry into the SON. Resting membrane potential was not different between sham and HF rats (sham: –63.3 ± 1.0 mV, HF: –62.7 ± 0.5 mV, n= 8 and 11, respectively, p= 0.6 unpaired t test). Optogenetic stimulation of SON_OT_-CeL fibers in sham rats resulted in a robust hyperpolarization (∼7 mV on average) and silencing of CeM neurons (p< 0.0001, paired t test). In line with a diminished release of OT described above, this effect was significantly smaller (∼2 mV on average) in CeM neurons of HF rats (p< 0.0001 vs sham, unpaired t-test) (**Figure 5B,C**, **Table 1**). A similar blunted inhibitory response in CeM neurons from HF rats was observed when we pharmacologically activated OTRs using the OTR agonist TGOT (1 µM, 3 min) (p< 0.0001 vs sham, unpaired t-test) (**Figure 5B,C**, **Table 1**). Together, these results support a blunted effect of OT in inhibiting output neurons from the CeM in rats with HF, an effect likely due to both presynaptic (i.e., diminished release of OT) and postsynaptic (i.e., diminished OTR expression) mechanisms.

**Figure 5.**
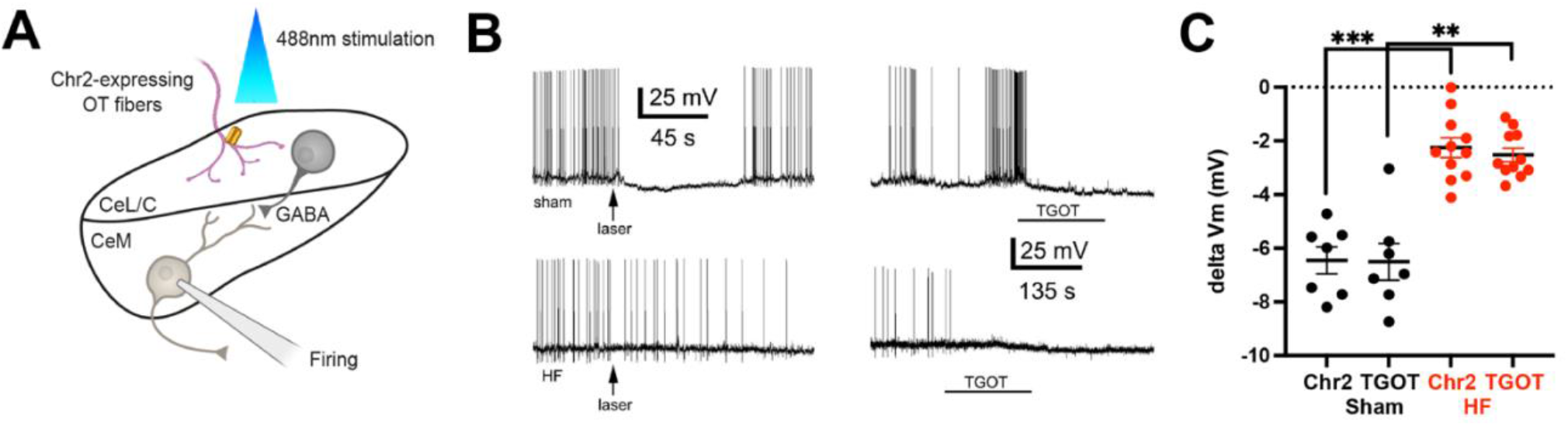
Electrophysiological assessment of CeL→CeM microcircuit functionality in HF rats. **A** Cartoon summarizing the experimental approach used. **B** Representative examples of inhibitory responses in CeM patched neurons evoked by laser stimulation (arrows, 488nm wavelength, 30s pulse) of Chr2-expressing OT fibers (left traces) or following bath application of 1 µM TGOT (right traces) in a sham rat (upper traces) or a HF rat (lower traces). **C** Summary data showing changes in membrane potential (delta Vm) evoked by either optogenetic stimulation (Chr2) or TGOT in CeM patched neurons in sham (black) and HF (red) rats. ***p< 0.001 and **p< 0.01, Mann-Whitney test.

**Table 1.**
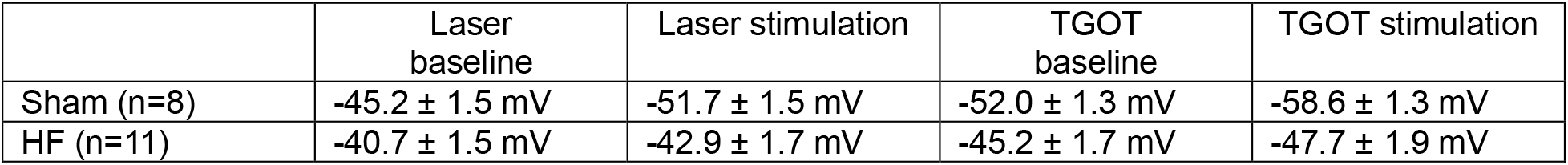
– Summary of membrane potentials (Vm) before and after laser stimulation of Chr2 fiber or following bath application of the OTR agonist TGOT in CeM neurons of sham and HF rats.

|  | Laser baseline | Laser stimulation | TGOT baseline | TGOT stimulation |
| --- | --- | --- | --- | --- |
| Sham (n=8) | -45.2 ± 1.5 mV | -51.7 ± 1.5 mV | -52.0 ± 1.3 mV | -58.6 ± 1.3 mV |
| HF (n=11) | -40.7 ± 1.5 mV | -42.9 ± 1.7 mV | -45.2 ± 1.7 mV | -47.7 ± 1.9 mV |

### Blunted efficacy of OT in driving GABAergic activity in CeM output neurons in rats with HF

To determine whether the blunted effect of OT on CeM output neurons in HF rats was due to a blunted effect of OT on GABAergic neuron activity (i.e., OTR-expressing neurons in the CeL) (**Figure 6A**), we monitored GABAergic IPSCs in CeM neurons using a high chloride internal solution before and after activation of OTRs after application of TGOT. As shown in **Figure 6B and C**, TGOT induced a robust increase in frequency as well as amplitude of GABAergic IPSCs in CeM neurons of Sham rats (∼150% and 100% respectively, p< 0.05 for both parameters, Wilcoxon test). Conversely, an almost completely blunted effect was observed in CeM neurons from HF rats (p< 0.01 vs. sham for both IPSC frequency and amplitude, Mann-Whitney test, **Figure 6C**, **Table 2**). While a tendency for higher frequency and amplitude of IPSCs was observed in HF rats, differences were not statistically significant.

**Figure 6.**
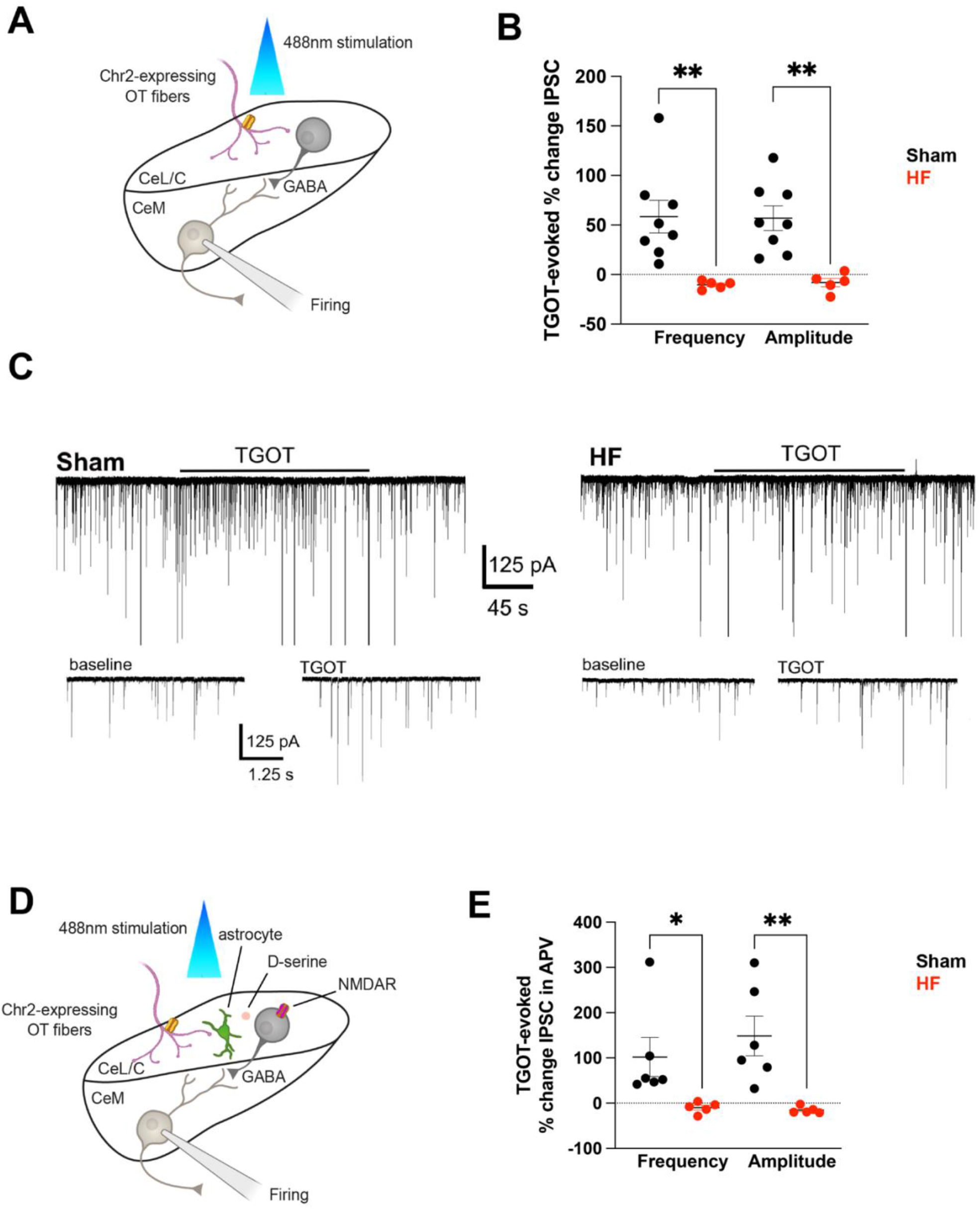
Analysis of TGOT-evoked iPSCS in the CeM of HF rats. **A** Cartoon summarizing the experimental approach used in B and C. **B** Summary data showing changes in IPSC frequency and amplitude evoked by TGOT in sham (black) and HF (red) rats. **C** Representative examples of voltage clamp traces recorded in CeM neurons showing IPSCs (inward currents) before and after TGOT application (arrow, 1 µM 3 min) in a sham (left) and a HF (right) rats. The lower traces show expanded segments at baseline and TGOT to better display individual IPSCs. **D** Cartoon summarizing the experimental approach for data shown in E. **E** Summary data showing changes in IPSC frequency and amplitude evoked by TGOT in slices preincubated with the NMDA receptor blocker APV (100 µM) in sham (black) and HF (red) rats. *p< 0.05, **p< 0.01, Mann-Whitney test.

**Table 2.** – Summary GABAergic inhibitory postsynaptic current (IPSC) properties before and after bath application of the OTR agonist TGOT in CeM neurons of sham and HF rats.

|  | IPSC frequency baseline | IPSC frequency TGOT | IPSC amplitude baseline | IPSC amplitude TGOT |
| --- | --- | --- | --- | --- |
| Sham (n=6) | 0.77 ± 0.2 Hz | 1.58 ± 0.30 Hz | 57.34 ± 11.6 pA | 116.6 ± 31.2 pA |
| HF (n=5) | 1.85 ± 0.5 Hz | 1.63 ± 0.52 Hz | 86.0 ± 10.0 pA | 75.5 ± 5.7 pA |

All IPSC currents were blocked by the GABA_A_ receptor blocker picrotoxin (400 µM, n=3, data not shown), further supporting the identity of the recorded IPSCs. These results support a blunted OT-driven CeL➜CeM GABAergic transmission in HF rats.

We recently showed that activation of OTRs in astrocytes also contributes to OTR-mediated activation of CeL neurons and inhibition of output CeM neurons ^30^, an effect thought to be mediated by astrocytic release of D-serine and subsequent activation of glutamate NMDARs in CeL neurons ^30^. Given this, and our results showing a diminished number of astrocytes expressing OTRs in rats with HF (**Figure 4**), we aimed to investigate the contribution of CeL neuroglial interactions to the blunted effect of OT in HF rats. To block OTR-astrocyte-CeL neuroglial pathway, we repeated experiments in the presence of an NMDAR blocker (APV, 100 µM) (**Figure 6D,E**) to interfere with the astrocyte-to-CeL GABA neuron communication ^30^. We found that in sham rats, TGOT still induced a significant increase in both the frequency and amplitude of IPSCs (p< 0.01 for both parameters, Wilcoxon test), although the magnitude of the effect was considerably less than that evoked in control conditions (i.e., 100% vs 50% in control and APV conditions, respectively). As before, an almost complete blunted effect of TGOT in the presence of APV was still observed in CeM neurons from HF rats (p< 0.01 vs sham for both IPSC frequency and amplitude, Mann-Whitney test, **Figure 6C**, **Table 3**). Together, and as we previously showed ^30^, these results indicate that in Sham rats, astrocytes contribute to OT effects on CeL➜CeM GABAergic transmission. However, since the differences between sham and HF persisted even in the presence of the NMDAR blocker APV, we interpreted these results as indicative that OT signaling in astrocytes does not constitute a major contributor to blunted OT effects on CeM➜CeL GABAergic transmission.

**Table 3.**
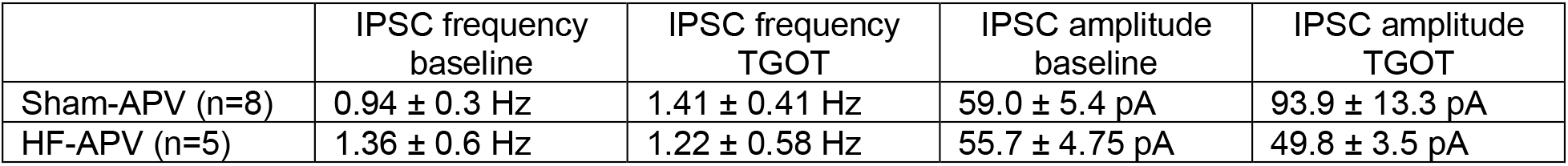
– Summary GABAergic inhibitory postsynaptic current (IPSC) properties before and after bath application of the OTR agonist TGOT in the presence of the NMDA receptor blocker APV in CeM neurons of sham and HF rats.

## Discussion

In this study, we combined a diverse array of electrophysiological, anatomical, viral, pharmacological and imaging approaches to study an OT-sensitive CeL➜CeM GABAergic circuit in the rat ischemic HF model. The critical role of GABAergic (OTergic) signaling within the CeA has been demonstrated multiple times before and the CeL➜CeM circuit has been shown to contribute to anxiety-like behavior or depression in several studies including work from our lab ^30,31,33^ and others ^48–52^. The main findings of our study include: i) no change in the number or electrophysiological properties of CeA-projecting SON OT neurons, which provides the main source of OT to this circuit, ii) a diminished OT cargo content along with a blunted activity-dependent release of OT from CeM axonal terminals from OTergic terminals in the CeM, iii) alterations in neuronal and astrocytic OTR expression within the CeA and iv) blunted OT-mediated modulation of astrocyte-independent CeM➜CeL GABAergic transmission, in the rat ischemic HF model. To the best of our knowledge, our study is the first of its kind reporting changes to an OTergic-GABA microcircuit that is involved in mood and memory regulation in a cardiovascular disease model.

### The OT-sensitive CeL**➜**CeM pathway and its role in mood regulation

The GABAergic microcircuit connecting the CeL to the CeM has garnered significance in the regulation of mood and memory ^48,53–57^. Prior research has underscored the crucial role of the hypothalamic OT pathway in modulating this intricate circuitry ^30,31^. Already in 2005, Huber and colleagues demonstrated the existence of an anatomically confined, OT-sensitive subpopulation of OTR-expressing neurons within the CeL^38^. Notably, emerging evidence has linked alterations in mood regulation, encompassing conditions like anxiety and depression, to prevalent cardiovascular diseases including hypertension^58–61^ and HF^8,10,11,39,62–64^. Pertinently, disruptions in OT signaling have been previously documented in various HF models ^23,26,40^. Building upon these findings, we put forth the hypothesis that perturbations in the OT-sensitive CeL➜CeM GABAergic pathway could potentially serve as a contributing substrate to the development of mood disorders within the context of heart failure. When studying perturbations to OT signaling in HF, three major aspects need to be considered: i) the source of OT, ii) the involved OT-ergic pathway and iii) the target structures and involved cell types, which we discuss below.

### Source of OT in the CeA

Previous studies have established the SON and PVN as the primary sources of OT synthesis within the brain ^33,65–67^. Our investigation employed a comprehensive approach, combining viral tracing and nucleus-specific anatomical analyses, which reaffirmed the SON’s role as the principal contributor of OT to the CeL (**Figure 1a-c**). In the context of our present study, meticulous examination of the CeA-projecting OTergic neurons revealed an absence of any alterations in the number of OT neurons in HF rats (**Figure 1d-f**). Further augmenting these findings, our employment of pharmacological techniques in tandem with electrophysiological assessments directly evaluated the excitability of these neurons (**Figures 1, 5 and 6**). These studies indicated that the fundamental intrinsic properties of these neurons, along with their repetitive firing patterns, remained unaltered, thereby corroborating the absence of changes in their excitability (**Figure 1g-i**). However, it’s worth acknowledging some limitations inherent to slice preparation, including the removal of peripheral inputs, and the absence of physiological variables. Yet, our prior utilization of slice preparations ^21,22,68,69^, coupled with our animal model ^39^, lends support to the notion that maladaptive plastic changes can indeed be translated from *in vivo* to *ex vivo* settings.

### OTergic SON➜CeA pathway

Our anatomical investigations have revealed a notable decrease in OT content within the axonal terminals connecting the SON to the CeL (**Figure 2**). However, our findings negate the possibility that this decline in OT immunoreactivity arises from a reduction in the number of terminals, as our results using AAV_OT_p_-Venus consistently demonstrate a similar fiber count between sham and HF rats (**Figure 2**). While intriguing, these results become even more perplexing in light of the absence of alterations in OT expression/content within the somata of SON-CeL neurons (**Figure 1**). One potential interpretation could be that the transport of OT to the terminals is compromised. Alternatively, these outcomes might signify the depletion of OT content due to heightened terminal activity during early HF stages ^23,24^. Clearly, further experiments are warranted to unravel the precise underlying mechanism responsible for the diminished OT content at the terminals. Irrespective of these potential mechanisms, these anatomical findings hold functional significance, as evidenced by our observation of a dampened activity-dependent release of OT (**Figure 3**). This was convincingly illustrated through our combined employment of optogenetic techniques and sniffer cell analysis, which unveiled reduced frequency, magnitude, and delayed responses of OT-sniffer cells following optical and selective activation of SON➜CeL OTergic fibers (**Figure 3**). Notably, the utilization of sniffer cells has been efficiently implemented to monitor both axon terminal and dendritic release of OT ^70,71^ and VP ^46^, boasting high sensitivity (nM detection) and spatiotemporal resolution, rendering them valuable tools for detecting endogenous neuropeptide release in the brain. Importantly, this approach has previously discerned differences in brainstem OT signaling in a different model of HF ^72^. In sum, our comprehensive anatomical and functional studies converge to support the notion of diminished OT levels in SON-CeL terminals and impaired activity-dependent neuropeptide release in the context of HF rats.

### Cellular targets of OT in the CeA

Our investigation delves into the multifaceted impact of OT on the CeL→CeM GABAergic microcircuit and its alterations in HF. Our findings go beyond compromised activity-dependent OT release from afferent terminals into the CeA, extending to a significant reduction in OTR expression. Anatomical changes in OTR expression were evident in both neurons and astrocytes, as we report a dichotomy of HF-induced changes on neuronal and astrocytic OTR expression in the CeA: While the percentage of OTR-positive neurons was not affected, we observed a ≍45% reduction of OTR expression levels (**Figure 4**). On the other hand, we observed a ≍35% reduction in OTR-positive astrocytes, but individual expression levels seemed unaffected (**Figure 4**). Intriguingly, only neuronal OTR downregulation was found to be physiologically relevant in our experimental context (**Figure 5**). To elucidate the functional implications of these molecular shifts, patch clamp recordings from CeM neurons were conducted. These neurons receive inputs from OTR-expressing CeL GABAergic neurons, known to be influenced by OT ^30,31,33,38^. As anticipated, in sham rats, both endogenous OT and the OTR agonist substantially suppressed CeM neuron firing activity. In stark contrast, HF rats exhibited a blunted response to both manipulations, underscoring the combined effect of reduced presynaptic OT release and diminished OTR expression on the CeL→CeM microcircuit. To further dissect this phenomenon, we monitored GABAergic synaptic activity in CeM neurons through direct measurement of GABA-mediated inhibitory postsynaptic currents (IPSCs) in voltage-clamp mode. In sham rats, OTR activation with TGOT led to increased frequency and amplitude of spontaneous IPSCs, effects largely absent in HF rats. This corroborated an overall compromised action of OT on the CeL→CeM GABAergic microcircuit in HF. Interestingly, we found a tendency for a higher basal GABAergic activity in CeM neurons in HF rats. This could seem in principle counterintuitive to the results described above. However, spontaneous activity represents all the GABAergic inputs into these neurons, not specifically those originating from OT-sensitive CeL neurons. Thus, it remains unclear at present whether changes in the basal degree of activity of CeL GABAergic neurons contribute to this effect. Nonetheless, our results compellingly show a blunted activation of the CeL➜ CeM inhibitory circuit in response to OTR activation.

Building on previous findings regarding the role of astrocytes in OT-mediated GABA function regulation, our study also revealed reduced OTR expression in these glial cells. We recently described the existence of an OT-sensitive astrocyte population within the CeL, which forms an astrocyte syncytium with neighboring astrocytes through gap junctions, and releases the gliotransmitter d-serine to activate nearby neurons ^30^. Thus, to ascertain the involvement of astrocytes, we repeated experiments in the presence of the NMDAR blocker APV ^30^. Interestingly, while the effect of the OT agonist on GABAergic activity was diminished in sham rats, the blunted OT effect in HF persisted, indicating a limited contribution of astrocytes to the observed modulation in the CeL→CeM GABAergic circuit. This finding suggests that the HF-induced changes in transmission are a result of impaired OT signaling through neurons, but not astrocytes. In line with this, we recently proposed that OT release within the CeA might inhibit neuronal Na^+^-activated K^+^ leak channels and that the coordinated dual activation of both NMDA and OTR could result in a potentiated neural response ^73^. In sum, our study presents a comprehensive view of the intricate OT-mediated regulatory mechanisms in this circuit, revealing significant alterations in HF, and strongly indicate HF-induced pathological alterations to the OT-sensitive CeL→CeM GABAergic circuit. Our current working hypothesis of impaired OT signaling in HF rats is summarized in a schematic illustration **on Figure S3**.

### Study limitations and conclusions

Our study has clear limitations. In HF, a myriad of processes including neuroinflammation^16,39,41^, hypoxia ^39,74^ and elevated circulating levels of neuropeptides like AngII ^39,75^ occur, all of which could potentially contribute to the changes reported in this study. We recently demonstrated that microglial activation, neuroinflammation, neuronal deterioration and apoptosis in the hippocampus of HF rats is predominantly driven by microglial AngII signaling ^39^. While future studies are warranted to address the triggering factors in HF, our data suggests that there is no single underlying mechanism that drives the HF-induced changes to the CeL➜CeM GABAergic microcircuit, given the multiple and varied levels of compromised reported herein. Another important caveat is that we performed all of our experiments at the established stage of HF (5-7 weeks after surgery). We don’t know when exactly the observed alterations to the SON➜CeA OTergic circuit occur, whether they persist over long periods of time and whether they precede or follow the previously reported cognitive and mood changes^10,39^. While our study provide detailed anatomical and functional information about changes in the OTergic SON➜CeL circuit and mechanistic data regarding altered CeM➜CeL GABAergic transmission, they do not prove that this is the underlying mechanism of the previously reported cognitive and mood changes in HF rats^10,39^. Follow-up behavioral experiments to unequivocally demonstrate that the OTergic SON➜CeL circuit underlying these behavioral impairments in HF rats could include a combination of CAV2-Cre injections into the CeA paired with AAV_OTp-DIO-hM3Dq injections into the SON to activate CeA-projecting OT neurons via application of CNO in a battery of behavioral tests. Nonetheless, the findings of this study suggest that targeting either CeA OTergic or GABAergic circuits pharmacologically could be feasible approaches for therapeutic interventions in HF patients suffering from cognitive decline, depression or mood changes.

Taken together, our study offers an initial glimpse into potential mechanisms underlying mood disorders in common cardiovascular conditions such as HF, although the precise link between the altered CeM➜CeL circuitry and depression and anxiety remains elusive. These findings open avenues for potential therapeutic interventions by enhancing either OT signaling or GABAergic function. The surge in research on OT agonists ^76–78^, including administration through intranasal routes ^79–81^, holds promise for clinical applications, although controversies persist ^79,82,83^. Importantly, the realm of GABA function improvement already benefits from several FDA-approved methods such as Zuranolone for post-partum depression ^84^, providing a tangible pathway for intervention. Albeit still controversial ^79,82,85^, targeting the OT system to ameliorate behavioral symptoms associated with various neuropsychiatric diseases has yielded encouraging results ^79,86^ that might have the potential to improve quality of life of affected individuals.

## Methods

### Animals

We used male Wistar rats (5-7 weeks old at surgery, 180-200g, Envigo, Indianapolis, IN, USA) for all experiments (total n=93). All experiments were approved and carried out in agreement to the Georgia State University Institutional Animal Care and Use Committee (IACUC) guidelines. To be consistent with our previous studies on heart failure, we only included male rats in the current study. Rats were housed in cages (2 per cage) under constant temperature (22 ± 2°C) and humidity (55 ± 5%) on a 12-h light cycle (lights on: 08:00-20:00). For immunohistochemistry and RNAScope experimental procedures, rats were killed by overdose of sodium pentobarbital and transcardially perfused with 0.1M phosphate buffer (pH 7.4) followed by 4% paraformaldehyde in phosphate buffer (pH 7.4). Brains were removed and post-fixed for 24 hours in 4% paraformaldehyde at 4°C followed by subsequent changes in escalating concentrations of sucrose in PBS (10, 20, 30% for 24 hours each).

### Heart failure surgery and echocardiography

To induce HF in rats, coronary artery ligation surgery was performed as previously described ^39^. Animals were anaesthetized using 4% isoflurane and intubated for mechanical ventilation until the end of the surgery. To exteriorize the heart, we performed a left thoracotomy. The ligation was performed on the main diagonal branch of the left anterior descending coronary artery. Animals received buprenorphine SR-LAB (0.5 mg/kg, S.C.; ZooPharm, Windsor, CO, USA) before the surgical procedure to minimize postsurgical pain. Sham animals underwent the same procedure except the occlusion of the left coronary artery. Four to five weeks after the surgery we performed transthoracic echocardiography (Vevo 3100 systems; Visual Sonics, Toronto, ON; Canada) under light isoflurane (2-3%) anesthesia to assess the ejection fraction (EF) and confirm the development of HF. We obtained the left ventricle internal diameter and the left diameter of the ventricle posterior and anterior walls in the short-axis motion imaging mode to calculate the EF. The myocardial infarct surgery typically results in a wide range of functional HF, as determined by the EF measurements. Rats with EF<40% were considered as HF (Sham 82.4±1.1%, HF 25.5±1.7%) (**See Figure S1** for additional echocardiographic parameters). Rats that underwent HF surgery but did not develop HF or displayed an EF>40% were not included in the study.

### Immunohistochemistry

Following pentobarbital-induced anesthesia (Euthasol, Virbac, ANADA #200-071, Fort Worth, TX, USA, Pentobarbital, 80mg/kgbw, i.p.), rats were first perfused at a speed of 20mL/min with 0.01M PBS (200mL, 4°C) through the left ventricle followed by 4% paraformaldehyde (PFA, in 0.3M PBS, 200mL, 4°C), while the right atrium was opened with an incision. Brains were post-fixated for 24 hours in 4% PFA at 4°C and transferred into a 30% sucrose solution (in 0.01M PBS) at 4°C for 3-4 days. For immunohistochemistry, 40 µm slices were cut using a Leica Cryostat (CM3050 S) and brain slices were kept in 0.01M PBS at 4°C until used for staining. Brain sections were collected in series using the merging anterior commissural as an initial landmark so that consistent collecting of brain tissue across all animals could be achieved. Brain slices were blocked with 5% Normal Horse Serum in 0.01M PBS for 1h at room temperature. After a 15-min washing in 0.01M PBS, brain slices were incubated for 24hrs (48hrs in case of PS38) in 0.01M PBS, 0.1% Triton-X, 0.04% NaN_3_ containing different antibodies: 1:1000 PS38 Neurophysin-I (gift from Harold Gainer, antibody now commercially available through Sigma-Aldrich MABN844), 1:1000 anti-glutamine synthetase (monoclonal mouse, Merck Milipore, MAB 302, clone GS-6) or 1:200 anti-oxytocin receptor (polyclonal rabbit, Alomone labs, AVR-013) at room temperature. Following 15-min washing in 0.01M PBS, sections were incubated in 0.01M PBS, 0.1% Triton-X, 0.04% NaN_3_ with 1:500 Alexa Fluor 488/594-conjugated donkey anti-rabbit/goat/mouse (Jackson ImmunoResearch, 711-585-152, 705-585-147, 715-545-151) for 4 hours at RT. Brain slices were washed again for 15 mins in 0.01M PBS and mounted using antifade mounting medium (Vectashield with DAPI, H-1200B).

### RNAScope

RNAScope reagents were purchased from acdbio (Multiplex v2 RNAScope kit and OTR rat probe). Opal 570 dyes were purchased from Perkin Elmer (Akoya Biosciences), Nuclease-free water and PBS were purchased from Fisher Scientific. Brains were processed as described under ***Immunohistochemistry*** using nuclease-free PBS, water, PBS and sucrose. We followed protocol as previously described ^39,40,47^.

### Analysis of OTR intensity, density, quantification and counting of cells and fibers

Density and intensity of OTR immunosignal were assessed in Fiji (NIH) using the threshold paradigm. Average pixel intensity was calculated using the Analyze<Measure function using collapsed 40µm sections (z-stack) images. For density analyses, raw images were opened and threshold was set individually for each image to achieve a near-perfect overlap of raw immunosignal and pixels used for quantifications. Optical density was then calculated via the Analyze<Measure function using the thresholded images so that only pixels above the threshold are included in the further image analysis. For quantifications we used at least 6-8 sections for each animal and brain region, if not indicated otherwise. Cells/neurons were counted manually using the cell counter plugin in Fiji. We used 8-12 sections containing dorsal CA2 per animal and sections were selected according to the indicated Bregma levels. OTergic fibers were counted manually in Fiji, while we defined a continuous line longer 2µm as a separate fiber. OTR immunoreactivity (OTR antibody) in CA2 astrocytes was assessed via co-localization analysis using Fiji as previously described ^30,87^. We considered astrocytes to be OTR-positive if at least 15% of GFAP immunoreactivity of each individual astrocyte overlapped with OTR protein immunoreactivity. To differentiate neurons from astrocytes for the OTR+ RNAScope experiments, we ran a separate set of experiments in which we immuohistochemically labeled for astrocytes (glutamine synthetase antibody) and neurons (NeuN antibody) along with DAPI nuclear staining. Using ImageJ algorithms, the nuclear size (DAPI area) of individually identified cell types was measured, and a frequency distribution histogram was built. As shown previously ^39^, neurons were readily differentiated from glial cells based on their significantly larger DAPI nuclear size. This approach allowed us to skip performing multiple additional immunostaining steps, which would have compromised the quality of our immuno-RNAscope experiments and subsequent mRNA probe signal intensity.

### Reverse transcription polymerase chain reaction (RT-PCR) and quantitative real time PCR (qPCR)

RNA extraction and isolation were performed using the miRNAeasy Mini kit (Qiagen, Cat. No. 217004) and the QIAzol Lysis Reagent (Qiagen, Mat. No. 1023537). 200µm-thick tissue sections were made in cryostat (–20°C, Leica, CM3050S) and punches from central amygdala (8-12 punches per animal) were collected and kept in dry ice until the RNA extraction procedure. RNA concentration was measured using NanoDrop One (Thermo Scientific) and was in the range of 260 – 330 ng/µl for all samples prior to cDNA synthesis. cDNA synthesis was performed using the iScript^TM^ gDNA Clear cDNA Synthesis Kit (BIO RAD, cat. no. 1725035) and the SimpliAmp Thermal Cycler (applied biosystems, Thermo Fisher Scientific) according to the manufacturer protocol. qPCR was conducted using the following 10x QuantiTect primers (diluted in 1.1 mL TE pH 8.0, final concentration: 200nM) purchased from Qiagen: OTR (QT100379036) and β-Actin (QT00193473). Housekeeping gene stability was assessed for each individual experiment using triplicate reactions. We only included datasets where the relative levels of β-Actin controls between sham and HF animals did not differ significantly. All individual qPCR reactions (brain region, primer and condition) were triplicated and averaged for further statistical analysis.

### Stereotaxic injection of viral vectors

Injection of viral vectors into the rat brain was performed in analogy to ^33^ using the Neurostar^TM^ injection robot. For viral infection of hypothalamic nuclei the following coordinates were used: SON (M-L ± 1.6mm, A-P – 1.4mm, D-V –9.0mm) and PVN (M-L ± 0.3mm, A-P –1.8mm, D-V –8.0mm). Injections into CeA were performed using the following coordinates: (M-L ± 4.5mm, A-P –2.5mm, D-V –8.0mm). Point of origin for the coordinates was Bregma and the Z level difference of Bregma and Lambda did not exceed 0.1mm ^88^. Injection volume per injection site was 300 nl, while all viruses used were in the range of 10^12^ –10^13^ genomic copies per ml. Retrograde labeling via CAV2-Cre and OT-promoter driven, cre-dependent viral expression was performed in analogy to ^31,40,44,45^. All experiments were performed after 4-5 weeks of viral expression.

### Quantitative analysis of OT release using Sniffer^OT^ cells

Full methods are previously described ^46^. Briefly, sniffer^OT^ cells were generated by culturing Chinese hamster ovary (CHO) cells in Dulbecco’s Modified Eagle Medium containing 10% w/v fetal bovine serum, 1% w/v penicillin–streptomycin, 1% w/v Na-Pyruvate, and 1% w/v NaCO3 filtered once through a Nalgene filtration system, and transfecting with pcDNA3.1+ containing human OT receptors cloned in EcoRI (5’) and XhoI (3’) (plasmid obtained from Missouri S&T cDNA Resource Center, Rolla, MO, USA) using lipofectamine, and stable overexpression was achieved by geneticin (500 mg ml^−1^) selection. 16-20 hours before experiment, Sniffer^OT^ cells were transiently transfected to express the red fluorescent genetically encoded calcium indicator R-GECO (GenScript, Piscataway, NJ, USA) with Fugene HD reagent (Promega, Madison, WI, USA). Sniffer^OT^ cells were resuspended in standard aCSF (in mM): 119 NaCl, 2.5 KCl, 1 MgSO4, 26 NaHCO3, 1.25 NaH2PO4, 20 D-glucose, 0.4 ascorbic acid, 2 CaCl2, and 2 pyruvic acid; pH 7.3; 300-305 mOsm) with trypsin (0.05%). Sniffer^OT^ cells were transferred via pipette directly onto the medial sector of central amygdala (CeM) of coronal brain slices submerged in aCSF in a perfusion-capable chamber of an upright microscope. After allowing Sniffer^OT^ 5 min to adhere to the slice, aCSF superfusion was resumed for 5 min to wash off any unattached Sniffer^OT^ from the slice before proceeding with imaging. Sniffer^OT^ cells adopted a rounded appearance when transferred to aCSF and no further overt morphological changes were observed during the equilibration period on the brain slices. Experiments were restricted to preparations that had at least five fluorescently visible sniffer cells in the field (∼15 on average). To record the calcium-induced fluorescence changes of Sniffer^OT^ cells, images were taken with the Dragonfly 200 confocal scanning system and iXON EMCCD camera (Andor Systems, Belfast, UK), at a rate of 1 Hz using piezo-driven z-series to maximize Sniffer^OT^ count per trial. The sniffer^OT^ cell fluorescence was imaged under a 561 nm excitation and the R-GECO calcium response was measured at 600 nm. During detection of OT release from terminals, slices were constantly superfused with aCSF at 32°C at ∼2 mL/min. Axon terminals containing AAV_OT_p_-Chr2mCherry were stimulated using an Andor Mosaic at 488 nm with 25 ms pulses at 8Hz. Imaging data were analyzed using ImageJ software (NIH). All data was background subtracted. For quantitative measurements, fractional fluorescence (F/F_0_) was determined by dividing the fluorescence intensity (F) within a region of interest by a baseline fluorescence value (F_0_) determined from 30 frames before stimulation. Peak calcium amplitude was the maximum F/F_0_ achieved after Chr2 stimulation. Latency (s) to the response was determined as the time between the end of the Chr2 stimulation and the start of the calcium response of each sniffer cell. Response duration (s) was the duration from the start of the response to the return baseline. The area under the curve (F/F_0_*s) was calculated using the integrated area for the duration of the response. Response rates are the number of cells that responded to both optogenetic Chr2 stimulation and an OT bolus divided by all cells that responded to the OT bolus. The OT bolus was delivered through the perfusion line via syringe (1 mL 0.1 µM; 0.1 mL s^-1^). Sniffer^OT^ cells that showed intrinsic oscillatory calcium activity were excluded from the analysis. To better display changes in fluorescence levels, images were pseudocolored using ImageJ.

### Patch-clamp electrophysiology in ex-vivo slices

Conventional patch-clamp recordings from neurons in the CeM were obtained from acutely obtained coronal slices from Sham and HF rats as previously described^89^. Extended optic tract and anatomical location of the stria terminalis were used as landmarks to identify the CeL region. These neurons were typically surrounded by mCherry-labeled, Chr2 containing fibers. Briefly, whole-cell current clamp recordings were obtained with an Axopatch 200B amplifier (Axon Instruments, Foster City, CA) and digitized using an Axon 1440B Digitizer (Axon Instruments, Foster City, CA). For current-clamp recordings, a conventional potassium gluconate-based internal solution was used. A similar approach was used to record from identified SON_OT_-CeL-projecting neurons. Input/output plots were generated from square pulses of 0pA to +180pA current injections in 20 pA increments, each lasting 1 sec. The spike frequency and counts were measured at each level and plotted as a function of the amount of current injected. For voltage-clamp recordings of GABAergic IPSCs, a high-chloride internal solution was used (in which potassium gluconate was replaced with KCl) to better detect IPSCs, as previously described ^90^. In this condition, and at the holding potential at which neurons were clamped (–70 mV), IPSCs appeared as large inward currents. IPSCs were detected and analyzed using MiniAnalysis. A detection threshold was set at −35 pA for IPSC peak, to extract IPSCs without contamination with glutamate-mediated excitatory postsynaptic current ^90^.. Is some experiments, as indicated in the text, axon terminals in the CeL containing AAV_OT_p_-Chr2mCherry were stimulated using an Andor Mosaic at 488 nm).

### Statistical analysis

All statistical analyses were performed using GraphPad Prism 8 (GraphPad Software, California, USA). The significance of differences was determined using two-tailed non-parametric tests (either unpaired (Mann-Whitney) or paired (Wilcoxon), one-way or two-way ANOVA or one-sample t-test, as indicated in the respective figure legends. Results are expressed as mean ± standard error of the mean (SEM). Results were considered statistically significant if p<0.05 and are presented as * for p<0.05, ** for p<0.01 and *** for p<0.0001 in the respective Figures.

## FIGURE LEGENDS

**Figure S1.**
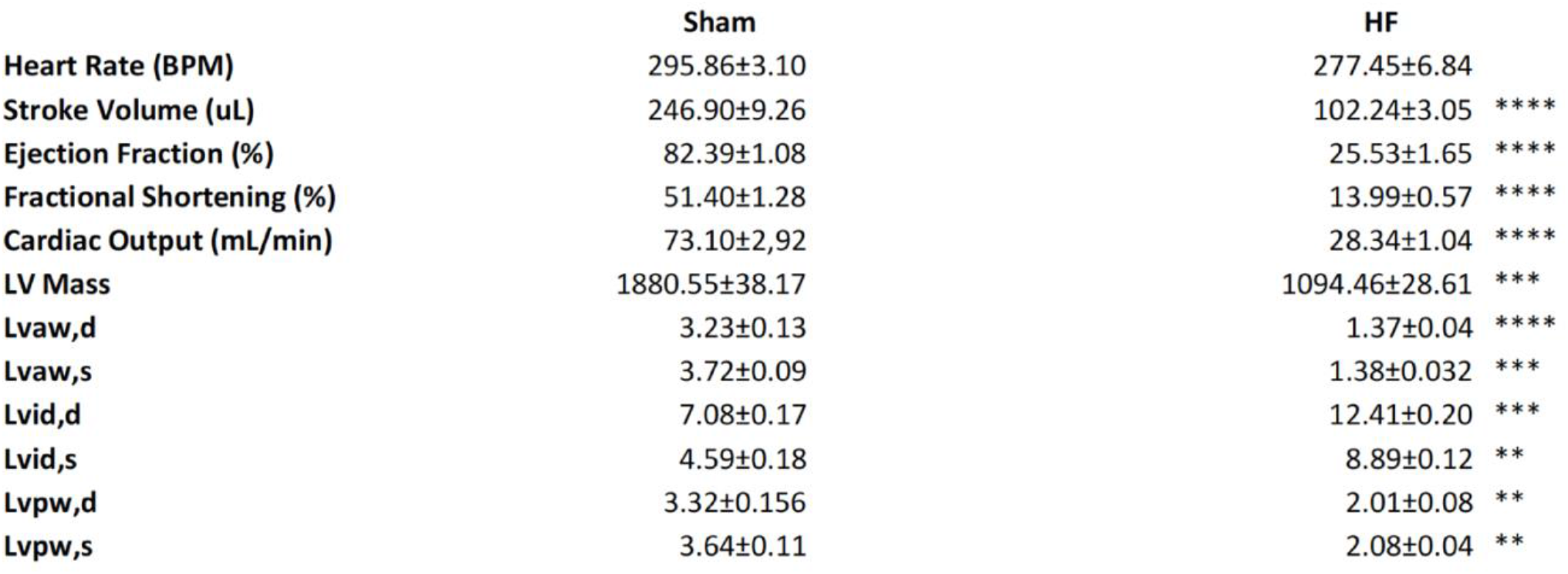
**A** Summary of assessed cardiac parameters in sham and HF rats via echocardiography. ***p<0.0001, ***p<0.001, **p<0.01 and *p>0.05 Mann-Whitney test.

**Figure S2.**
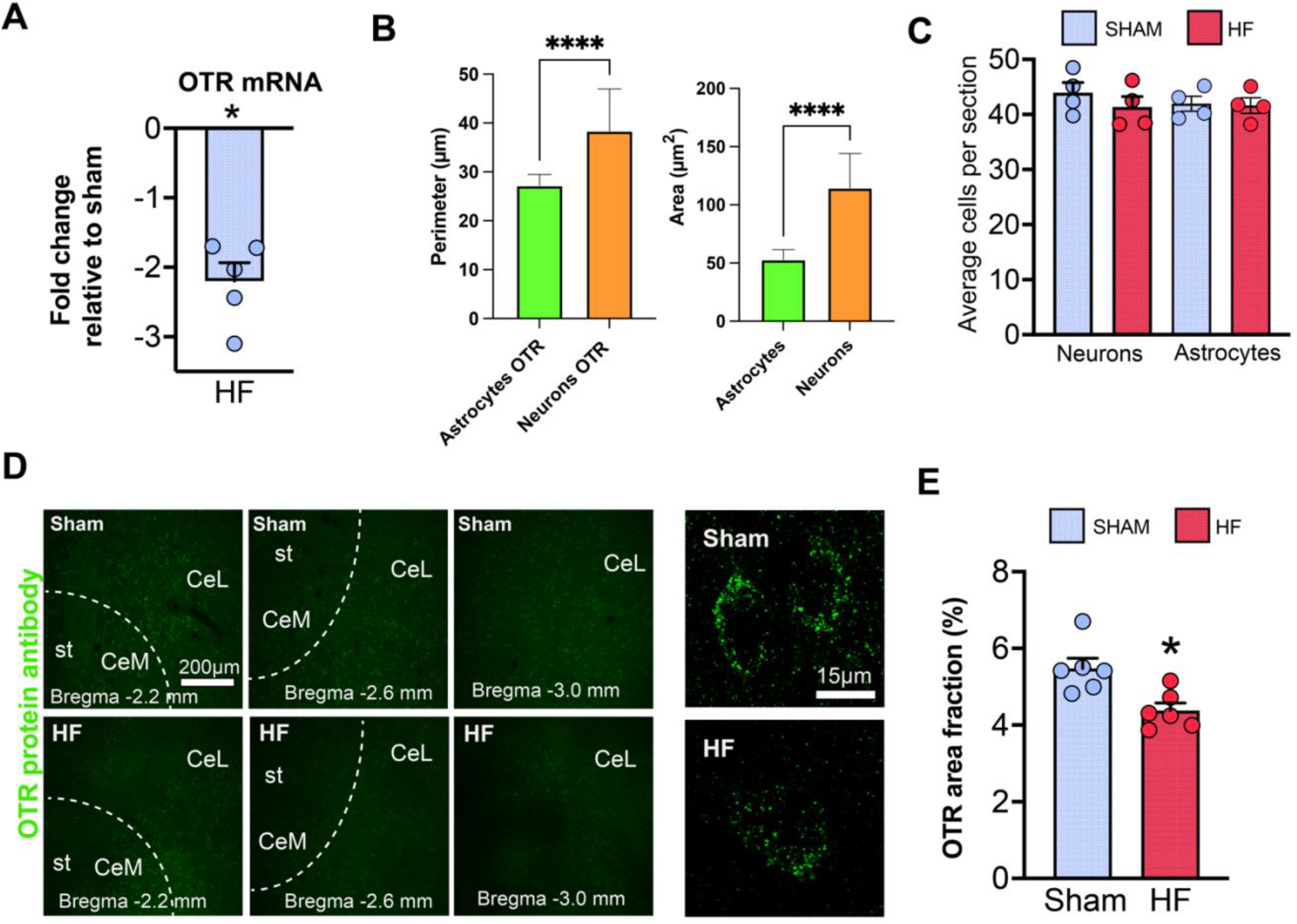
**A** CeA-specific tissue punches and qPCR for OTR mRNA in HF rats (n=5). **B** Fiji-based assessment of neuronal and astrocytic perimeter and area (sham and HF pooled). **C** Quantification of neurons and astrocytes within the CeA of sham and HF rats (n=4 per group). D Representative confocal images of CeA-containing brain sections stained with the OTR antibody from sham and HF rats at various Bregma levels, st=stria terminalis. High magnification images show OTR-positive puncta in individual cells. **E** Quantification of OTR area fraction within the CeA in sham and HF rats. *p<0.05 one-sample t-test and ****p<0.0001 Mann-Whitney test.

**Figure S3.**
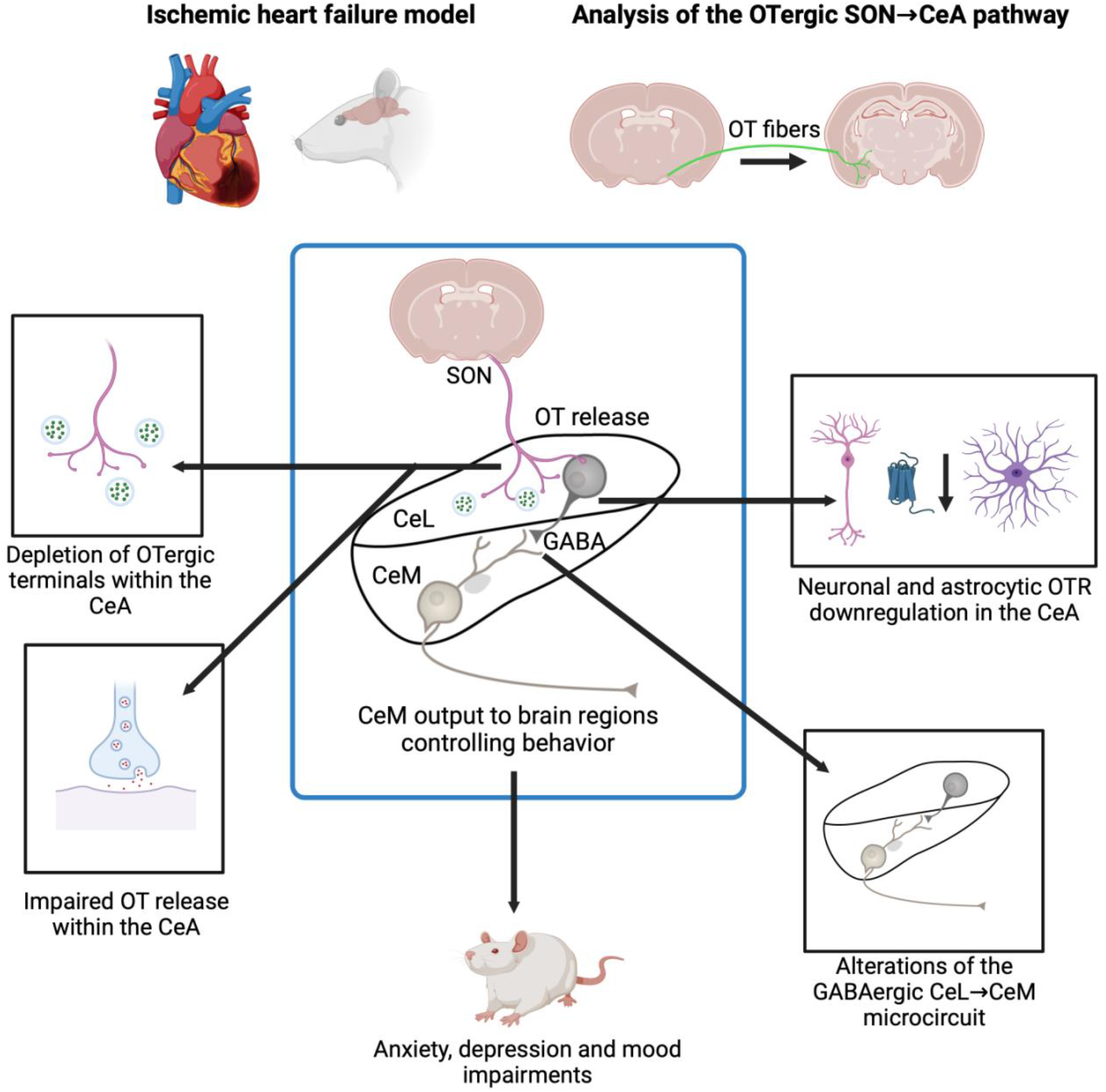
Summary of the current working hypothesis of impaired SON→CeA OT signaling in HF rats.

## ACKNOWLEDGEMENTS

Schematic illustrations on Figures 2, 5 and 6 were created using biorender.com

